# A Real-Time Functional Localization Method Based on Dynamic Connectivity for Coupled Brain Regions

**DOI:** 10.64898/2026.02.02.703208

**Authors:** Nils Jannik Heukamp, Vera Moliadze, Frauke Nees

## Abstract

In the rapidly growing field of dynamic connectivity (dyC)-based real-time functional magnetic resonance imaging (rtfMRI) neurofeedback training, participants receive feedback on the dynamical coupling of brain nodes based on real-time dynamic connectivity measurements between multiple regions-of-interest. Given the fundamental role of regions-of-interest in this context, their signal-to-noise-ratio is critical. In closely related activity-based rtfMRI neurofeedback, region-of-interest selection is guided by functional localization methods informed by the BOLD-signal during a functional localizer experiment conducted prior to neurofeedback. However, methods that account for dynamic coupling of multiple brain regions remain lacking.

Here, we develop a method based on dynamic connectivity between two brain areas that aligns with conventional activity-based localization approaches and is applicable for rtfMRI-NF.

The proposed pipeline adapts processing steps from activity-based methods and was successfully implemented in a dyC-informed neurofeedback study. We observed a significant increase in dyC signal-to-noise-ratio during the localizer experiment and the first three blocks of neurofeedback training. Importantly, dyC values also correlated with anxiodepressive symptom levels, serving as proof of sensitivity, as changes in dyC reflect not only task-related neural dynamics but also individual differences in anxiodepressive traits.

Our findings demonstrate that functional localization methods can be extended to dynamic connectivity, improving rtfMRI feedback accuracy, and with evidence to serve as sensitive measure for emotional states.

This approach may enable precise targeting of coupled brain regions in neurofeedback and holds potential for personalized clinical applications, with broader implications beyond neurofeedback discussed.

## Introduction

Dynamic connectivity-based rtfMRI neurofeedback targets modulation of specific aspects of functional brain networks (Fede et al., 2020; Hampson, 2021). In this approach, participants receive real-time feedback on the coupling of brain nodes derived from stochastic dependencies in blood-oxygen-level-dependent (BOLD) signals between multiple regions-of-interest (ROIs) (Goebel, 2021; Koush et al., 2013). Thus, ROI selection is critical, as it directly affects the accuracy of functional connectivity estimation and the subsequent neurofeedback signal (Goebel, 2021; Lührs et al., 2017).

This challenge encompasses not limited to dyC-based neurofeedback but also applies to activity-based fMRI neurofeedback, where functional localization approaches are commonly used (Cohen Kadosh et al., 2016; Fede et al., 2020; Goebel, 2021). In these approaches, targeted brain functions are deliberately elicited during a pre-training localizer experiment. Voxel-wise BOLD signals are analyzed using standard linear models, producing subject-specific functional maps that allow selection of ROIs with robust signal-to-noise-ratio (SNR) (Goebel, 2021; Lührs et al., 2017). These methods can be applied in (near) real-time, automated, and improve the quality of subsequent rtfMRI-measurements (Goebel, 2021; Lührs et al., 2017). However, no equivalent method exists for functional localization of dynamic coupling that accounts for dyC SNR across multiple brain regions.

To address this methodological gap, we developed a dyC-based functional localization method (dyC-MFL) for two brain regions, adhering to the principles of activity-based localization while remaining applicable in real-time neurofeedback. We describe the pipeline, challenges and solutions, and demonstrate its implementation in a dyC-based neurofeedback study. We further report results from both the localizer experiment and neurofeedback training and discuss implications and potential applications.

## Methods

### Pipeline development

To create a dyC-MFL pipeline compatible with activity-based pipelines from the literature (Goebel, 2021; Lührs et al., 2017), and adapted them for dyC, adding steps suitable for neurofeedback studies was then established. Details on pipeline development and technical setup are reported in the Results section.

### Neurofeedback Study

The dyC-MFL was implemented in a neurofeedback study targeting prefrontal-limbic coupling during pain processing, specifically the dyC between the dorsolateral prefrontal cortex (DLPFC) and amygdala. The study was registered with the German Clinical Trial Register (DRKS00024713) and approved by the Ethics Committee of the University Medical Center Schleswig Holstein (D 433/21). Four neurofeedback sessions were conducted on consecutive days. The first day included anatomical measurements, the localizer experiment, a five-minute resting-state run, and neurofeedback training. Analyses included 17 healthy participants with complete datasets.

### Subjects & Psychological Assessment

Participants were recruited via flyers and online advertisements. Inclusion criteria required age ≥18, no chronic pain, and no current or past psychological disorders except for prior remitted anxiety or affective disorders. Therefore, in addition to self-report, we performed the German version of the SCID-5 CV and in case of anomalies in the SCID-5 PQ also the SCID-5 PD in an on-site assessment prior to the NF-training (Beesdo-Baum et al., 2019). Psychological assessment included the Hospital Anxiety and Depression Scale (HADS, Snaith & Zigmond, 1995).

### Localizer Experiment Setup

The localizer aimed to elicit pain-related vs. pain-unrelated processing using a block design: six 50s blocks of electrical pain stimulation interleaved with seven 25s arithmetic tasks. Pain was induced using monopolar transcutaneous electric stimuli (Digitimer DS7A) targeting Aδ nociceptive fibers. Stimulation strength was individualized at 70% between pain threshold and pain tolerance. Arithmetic tasks involved continuous subtraction of 2 from randomly generated numbers between 100 and 500. The experimental procedure was implemented in PsychoPy (Peirce et al., 2019).

### Neurofeedback training

Each session included six runs alternating up- or downregulation, each with six blocks with active dyC feedback. Start condition and feedback direction were randomized. Analyses focus on the first run of the first session, to minimize neurofeedback-induced adaptation (see below).

### MRI Acquisition

MRI was acquired using a 3-Tesla Siemens MAGNETOM Cima.X MRI scanner. One T1-weighted SK\SP\MP single-echo structural MRI data was collected (208 slices; repetition time, TR=2200ms; echo time, TE=2.46ms; flip angle, FA=8°; field of view, FOV=230x230mm; matrix size=256x256; voxel size=0.9x0.9x0.9mm). For the localizer experiment segmented k-space single-echo fMRI data were collected (60 slices in interleaved ascending order; TR=1000ms; TE=38ms; FA=31°; FOV=204x204mm; matrix size=84x84; voxel size=2.43x2.43x2.4mm; MB factor=6). This was adopted from a previous study optimizing sequence for rtfMRI neurofeedback (Renz et al., 2023). The run duration was 8 minutes, during which 480 volumes were acquired.

#### Hardware

dyC-MFL computations were performed on a GIGABYTE AERO laptop (i7-11800H, 32 GB RAM, RTX 3070).

### Quantification of Outcomes and Statistical Analysis

For statistical analyses on differences and correlation we used two-sided t-tests with alpha=0.05.

#### Localizer Experiment

dyC SNR was estimated from p-values derived from GLM contrasts of pain vs. arithmetic blocks, corrected for multiple comparisons. ROIs derived from dyC-MFL were compared against template-based ROIs from atlases or meta-analyses.

#### Neurofeedback Training

SNR of dyC was quantified using first-run data, with block-wise and cumulative analyses. Associations between dyC and psychological measures (HADS) were also evaluated.

## Results

### Conceptual Development of dyC-MFL

We **adapted four key steps from activity-based MFL to dyC** (Figure 1, Supplemental Text 1):

**Figure 1.**
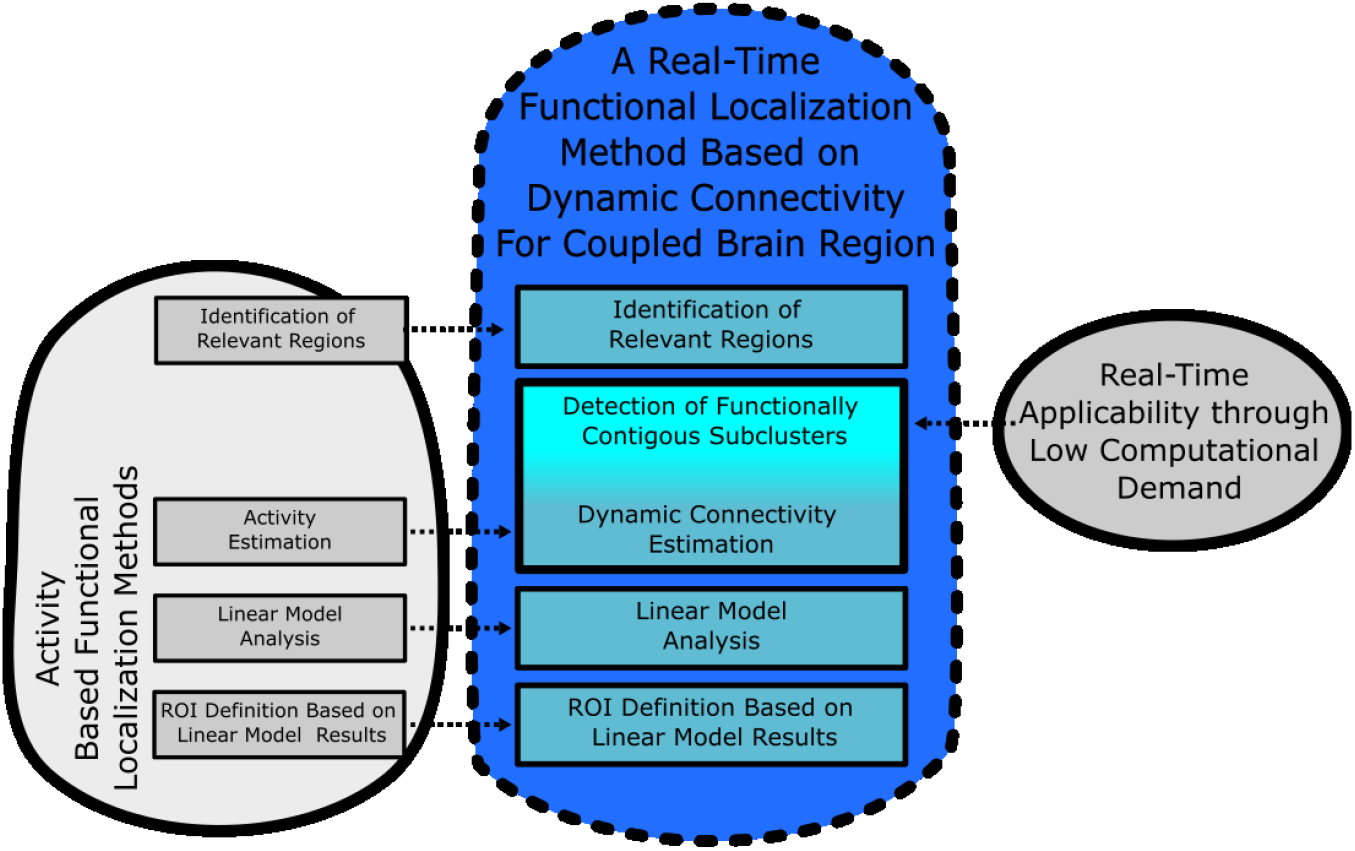
Schematic illustration of the developed dyC-based method of functional localization and its links to the objectives in the development. This were 1) to align it with existing activity-based procedures (left) and 2) maintain applicability for neurofeedback in (near) real-time (right). Left: Processing steps identified in activity-based MFL-pipelines. Steps that were transferable are colored dark blue. Additional substeps necessary for real-time applicability (right) are colored turquoise.

1. **ROI identification in MNI space**: extended to multiple brain areas.
2. **DyC estimation**: performed between **functionally contiguous subclusters** due to combinatorial explosion of voxel-wise dyC. Subclusters were defined via **k-means clustering of BOLD timeseries**.
3. **GLM analysis**: contrasts applied to dyC time series with minor adjustments for dyC-specific delays.
4. **ROI selection**: candidate ROI pairs with **highest dyC SNR** identified for neurofeedback.

### Technical Implementation

The pipeline successfully completed <5 minutes processing, compatible with the resting-state interval between localizer and neurofeedback.

#### Step 1: Identification and definition of relevant brain areas

First, definition masks for the amygdala and DLPFC were defined by thresholding meta-analytic z-maps in MNI space to the top 100 voxels. These maps were obtained from Neurosynth (Yarkoni et al., 2011) and reflect the meta-analytical association with the terms “DLPFC” or “Amygdala”, respectively (see Figure 2). The resulting masks were projected into individual subject space by co-registering each participant’s anatomical T1-weighted image to MNI space using TurboBrainVoyager (RRID: SCR_014175)(Goebel, 2012).

**Figure 2.**
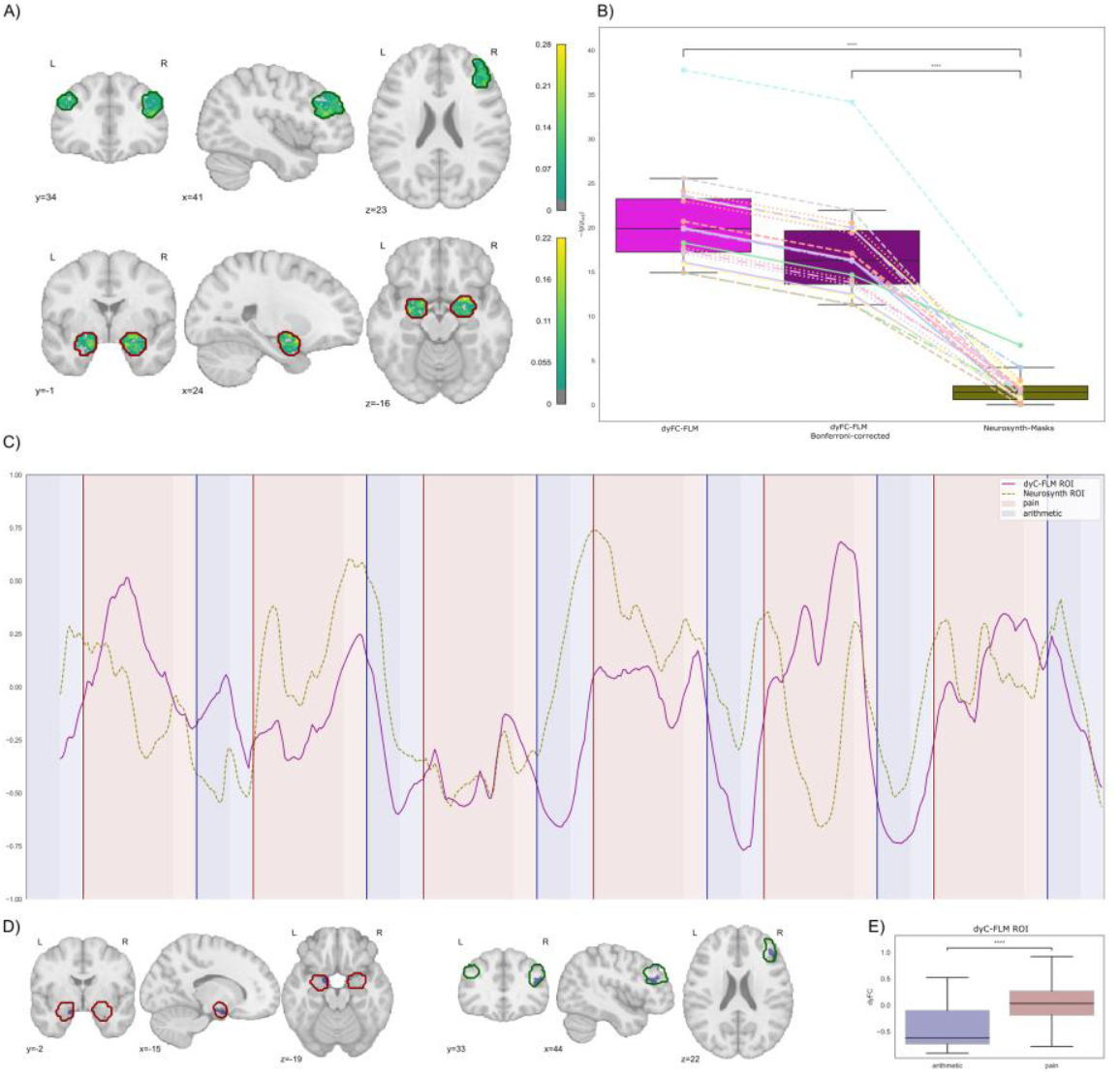
Outcomes of the Implementation in the neurofeedback study. (A)-(B): Results of the dyC-MFL across subjects (**A**) Relative Frequency of a voxel within the Neurosynth top 100 voxel masks assigned as voxel within the dyC-MFL optimized ROIs. Top DLPFC; Bottom amygdala. (**B**) Boxplots of the p-values (-log10) derived from the comparison of dyC between pain and arithmetic blocks. Left uncorrected, middle corrected for multiple comparisons (Bonferroni) during the dyC-MFL, respective values for the dyC between the Neurosynth top 100 voxel masks.. (**C**)-(**E**) Example results of a single subject run of dyC-MFL. (**C**) DyC estimation between the ROIs which were optimized by the dyC-MFL as compared to the dyC between the Neurosynth top 100 voxel masks. (**D**) dyC-MFL optimized amygdala (left), respectively DLPFC (right) ROI in blue within the respective borders of the Neurosynth masks. (**E**) Boxplot of the dyC of the selected ROI pair during the pain as compared to the arithmetic task. **** p<.0001.

#### Step 2: DyC Estimation Between Candidate Regions of Interest

Voxel-wise BOLD signals within the amygdala and DLPFC masks were extracted and denoised. rtfMRI preprocessing was performed using TurboBrainVoyager. The first five volumes of the localizer run were discarded to avoid saturation effects. Remaining images were 3D motion-corrected by alignment to the first volume and spatially smoothed (FWHM = 5 mm).

BOLD time series were extracted using the template masks and further denoised using nilearn’s signal.clean function, including six motion parameters, a linear drift term, and an intercept computed by TurboBrainVoyager (Goebel, 2012).

To identify candidate ROIs reflecting functionally contiguous subregions of the amygdala and DLPFC we applied a k-Means timeseries clustering algorithm implemented in TsLearn (Tavenard et al., 2020). This computationally efficient, data-driven approach is well suited for voxel-wise BOLD clustering when limited numbers of clusters is desired (Thirion et al., 2014). To allow for different levels of spatial granularity while constraining computational demand, the number of clusters incrementally increased until more than 4000 possible inter-regional cluster combinations were obtained . This threshold ensured that the total processing time of the dyC-MFL pipeline remained below five minutes, allowing completion during the intervening resting-state run.

Dynamic connectivity time series were then estimated for each pair of candidate ROIs spanning the amygdala and DLPFC. Consistent with previous work, dyC was computed using a sliding-window approach with a window length of 15 frames/ seconds (Zhao et al., 2019; Zich et al., 2020) . For each window, connectivity between the DLPFC and amygdala was estimated using partial correlation, controlling for a confounding physiological signal. This confound was defined as the mean BOLD signal from 100 randomly selected white-matter voxels, drawn from the TemplateFlow MNI152Asym white-matter template (Ciric et al., 2021) and located within the bounding box defined by the outermost voxels of the amygdala and DLPFC masks. All computations were performed using in-house scripts based on NumPy and Pandas (Harris et al., 2020; The pandas development team, 2020).

#### Step 3: General Linear Model Analysis

The step aimed to assess the SNR of each dyC timeseries with respect to pain processing. For this purpose, dyC values during pain blocks were contrasted against arithmetic blocks using general linear model, which in this two-condition design corresponds to a a two-sample t-test.

To reduce contamination by anticipatory processes, the final10 time points of each arithmetic block were excluded, as pain anticipation is known to jointly activate the amygdala and DLPFC (Mackiewicz et al., 2006; Nitschke et al., 2006). Likewise, the final 10 time points of each pain block were discarded to account for potential habituation effects (Bordi & LeDoux, 1992; Breiter et al., 1996). Statistical testing was implemented using SciPy (Virtanen et al., 2020).

#### Step 4: The Identification of the Regions of Interest

The final step aimed to identify the pair of candidate ROI masks exhibiting the strongest pain-specific dynamic coupling. Candidate ROI pairs were compared based on their SNR, indexed by the p-values obtained from GLM analysis. The ROI pair with the lowest p-value was selected and transformed back into MNI space for use in subsequent neurofeedback training runs and sessions using TurboBrainVoyager (Goebel, 2012).

### Implementation Outcomes

#### Localizer

dyC-MFL ROIs showed highly significant dyC differences between pain and arithmetic blocks (mean p<1.5×10^−16^). dyC SNR was significantly higher than template-based ROIs (t=18.77, p<.0001). Outcomes of the implementation in the localizer run and an exemplary dyC of a single subject are visualized in Figure 2 and ROI characteristics are provided in Table 1.

**Table 1.** Characteristics of ROIs derived by the dyC-MFL. Number of voxels and spatial overlap between subjects (number of voxels shared by two subjects proportional to their average number of voxels in the ROI).

|  | N(voxels) |  |  |  |  | % overlap |  |  |  |  |
| --- | --- | --- | --- | --- | --- | --- | --- | --- | --- | --- |
|  | mean | std | min | median | max | mean | std | min | median | max |
| **AMY** | 54.1176 | 21.6474 | 29 | 47.0 | 93 | 4.6139 | 11.5154 | 0 | 0 | 95.4545 |
| **DLPFC** | 56.4706 | 19.6345 | 27 | 53.0 | 97 | 5.3384 | 8.6587 | 0 | 2.0518 | 56.8966 |

AMY: amygdala, DLPFC: dorsolateral prefrontal cortex, std: standard deviation.

#### Neurofeedback Training

dyC SNR in the first and third blocks exceeded significance threshold (Table 2 & Figure 3). Cumulative analysis over three blocks also showed significance. dyC differences correlated with HADS total score (r=0.6577, p=0.006, Figure 3), robust to control for regulation condition.

**Table 2.** Results from SNR analyses in the first run of the neurofeedback training.

|  | block | mean | std | SEM | 95%-CI |  | t | p |
| --- | --- | --- | --- | --- | --- | --- | --- | --- |
|  |  |  |  |  | [LL, | UL] |  |  |
| single | 1 | 3.8533 | 3.1826 | 0.7956 | 2.1666 | 5.5400 | 3.2078 | 0.0055 |
|  | 2 | 3.6593 | 5.4188 | 1.3547 | 0.7875 | 6.5312 | 1.7408 | 0.1009 |
|  | 3 | 2.9460 | 2.9867 | 0.7467 | 1.3631 | 4.5289 | 2.2031 | 0.0426 |
|  | 4 | 1.7761 | 1.5145 | 0.3786 | 0.9734 | 2.5787 | 1.2546 | 0.2276 |
|  | 5 | 1.7323 | 1.8260 | 0.4565 | 0.7646 | 2.7001 | 0.9448 | 0.3588 |
|  | 6 | 3.6562 | 4.0432 | 1.0108 | 1.5134 | 5.7990 | 2.3300 | 0.0332 |
| cumulated | 1 | 3.9989 | 4.7738 | 1.1934 | 1.4689 | 6.5289 | 2.2606 | 0.0381 |
|  | 2 | 3.3050 | 3.0425 | 0.7606 | 1.6925 | 4.9174 | 2.6346 | 0.0180 |
|  | 3 | 4.2259 | 4.4557 | 1.1139 | 1.8645 | 6.5874 | 2.6258 | 0.0184 |
|  | 4 | 2.6856 | 2.8883 | 0.7221 | 1.1548 | 4.2163 | 1.9174 | 0.0732 |
|  | 5 | 2.1130 | 2.4066 | 0.6016 | 0.8375 | 3.3884 | 1.3495 | 0.1960 |
|  | 6 | 2.4484 | 3.0748 | 0.7687 | 0.8189 | 4.0780 | 1.4926 | 0.1550 |

Std: standard deviation, SEM: Standard Error of the Mean, CI: confidence interval, LL: lower limit, UL: upper limit

**Figure 3.**
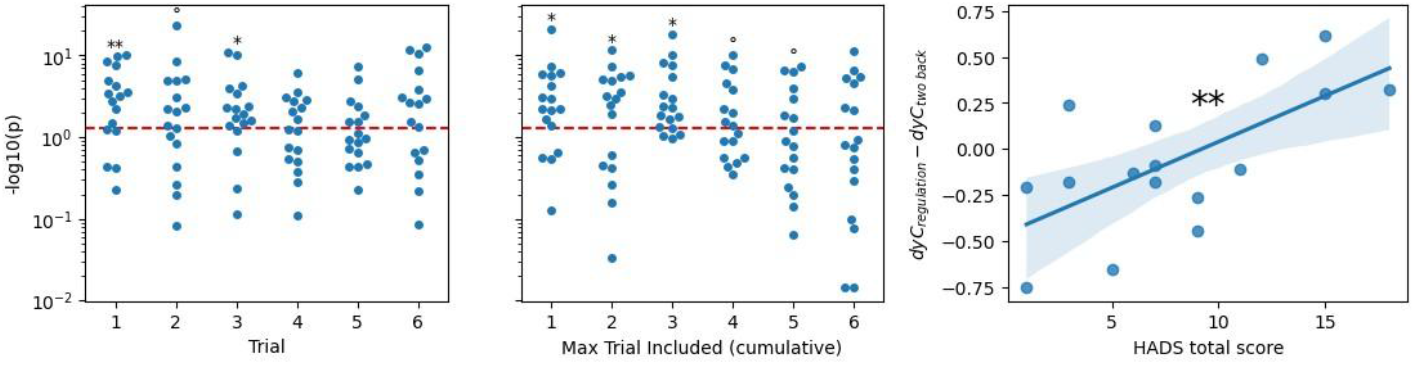
Outcomes of the MFL in the first neurofeedback training run. Left: negative decadic logarithmic p-values as indicator of SNR per trial in the first run with red line indicating alpha=0.05. Middle: negative decadic logarithmic p-values as indicator of SNR cumulated over trial in the first run with red line indicating alpha=0.05. Right: Association of dyC difference between regulate/pain and arithmetic in the cumulated first three blocks and the HADS total score. **p<.01, *p<.05, °p<.2.

## Discussion

We present a method (dyC-MFL) for functional localization of two coupled ROIs, aligned with activity-based approaches and implementable in real-time neurofeedback.The dyC-MFL enhances SNR, transfers from localizer to neurofeedback, and identifies ROIs associated with relevant psychological measures. This method extends functional localization to dynamic coupling, supporting personalized brain connectivity interventions.

While we implemented the dyC-MFL in a dyC-based neurofeedback study, previous dyC-based fMRI-neurofeedback studies also used a-MFL (Koush et al., 2017; Spetter et al., 2017; Zhao et al., 2019). dyC-MFL is suitable for neurofeedback targeting mutually interacting regions, while activity-based MFL remains valid when joint activity without mutual influence is the focus.

We further observed that dyC-signal derived from dyC-MFL during neurofeedback correlated significantly with psychometric measures of well-being, indicating potential sensitivity to psychological states. This suggests that, beyond improving SNR, dyC-MFL may enhance neurofeedback protocols by precisely targeting psychologically sensitive neural processes. These findings may may have clinical relevance, as numerous neurological and mental disorders are associated with altered brain connectivity alterations (Filippi et al., 2013; Fornito et al., 2015).

Beyond fMRI-neurofeedback, dyC-MFL has multiple potentials applications. Most prominently, non-invasive brain stimulation (NIBS) can be applied in closed-loop system (Bergmann, 2018; Mulyana et al., 2022; Soleimani et al., 2023) Whereas previous NIBS studies have defined ROIs based on static resting-state connectivity (Morriss et al., 2024), dyC-MFL may may allow targeted modulation of event-related brain functions, potentially enhancing closed-loop stimulation protocols.

Our study serves as a proof-of-concept. However, the effects of alternative dyC-MFL implementations and their methodological implications require further investigation. Further exploration of alternative clustering and dyC estimation algorithms could expand its utility.

In conclusion, we present a robust method for functionally localizing dynamic coupling between two brain areas, enhancing rtfMRI neurofeedback and laying the groundwork for personalized brain connectivity interventions.

## Data Availability

The authors confirm that the data supporting the findings of this study are available within the article.

