## Supplemental Text 1 for "A Real-Time Functional Localization Method Based on Dynamic Connectivity for Coupled Brain Regions"

The first step includes the identification of relevant brain areas and their definition in a standard space, typically MNI, and is fully transferable except that two or multiple brain areas are to be defined. In the second step, of activity-based MFL activity is estimated for every voxel within the defined mask of the relevant brain area (Goebel, 2021). In the transfer of this processing step a major challenge emerges. In contrast to BOLD-activity, dyC is defined as stochastic dependency of multiple BOLD-courses and thus cannot be estimated voxel-wise, but only between combinations of voxels. The number of specific combinations between voxels of multiple brain areas is exponential to the number of voxels contained in the masks. Calculating dyC for all combinations is given currently regular computational capacities precludes real time processing. We addressed this challenge by adding a step for detecting functionally contiguous brain area subclusters Based on results that machine learning-based dimensionality reduction of BOLD activity signals maintains connectivity signatures (Dadi et al., 2020). This can be based on any time-series algorithm and fine-tuned to the demands and computational capacities of individual technical set ups (see below for specific example). DyC is then estimated for every combination of functionally contiguous brain area subclusters. In the third step of activity-based MFL linear models are applied to the BOLD-series for estimating contrasts and respective SNR. Linear models can also be applied to dyC with minor changes. While contrasts in BOLD-activity models are corrected for hemodynamic delay, the delay in reactivity of dyC is not following hemodynamic response course but depends on the specific type of dyC measurement and thus needs to be addressed in respect to the individual study. Lastly, ROI are defined from the results of the linear mode. Here, again minor changes are necessary in the transfer from BOLD-activity to dyC. In BOLD-activity MFL thresholds are applied to define a cluster of voxels above a certain significance / SNR threshold. Given that in dyC-MFL there linear models were computed cluster combination- not voxel-wise the clusters from the cluster combination with the highest significance / SNR are to be defined as ROIs.
